# Genome-wide association study reveals candidate genes for flowering time in cowpea (*Vigna unguiculata* [L.] Walp)

**DOI:** 10.1101/2021.04.01.438123

**Authors:** Dev Paudel, Rocheteau Dareus, Julia Rosenwald, María Muñoz-Amatriaín, Esteban F. Rios

## Abstract

Cowpea (*Vigna unguiculata* [L.] Walp., diploid, 2*n* = 22) is a major crop used as a protein source for human consumption as well as a quality feed for livestock. It is drought and heat tolerant and has been bred to develop varieties that are resilient to changing climates. Plant adaptation to new climates and their yield are strongly affected by flowering time. Therefore, understanding the genetic basis of flowering time is critical to advance cowpea breeding. The aim of this study was to perform genome-wide association studies (GWAS) to identify marker trait associations for flowering time in cowpea using single nucleotide polymorphism (SNP) markers. A total of 367 accessions from a cowpea minicore collection were evaluated in Ft. Collins, CO in 2019 and 2020, and 292 accessions were evaluated in Citra, FL in 2018. These accessions were genotyped using the Cowpea iSelect Consortium Array that contained 51,128 SNPs. GWAS revealed seven reliable SNPs for flowering time that explained 8-12% of the phenotypic variance. Candidate genes including *FT, GI, CRY2, LSH3, UGT87A2, LIF2*, and *HTA9* that are associated with flowering time were identified for the significant SNP markers. Further efforts to validate these loci will help to understand their role in flowering time in cowpea, and it could facilitate the transfer of some of this knowledge to other closely related legume species.

## 1 Introduction

Cowpea (*Vigna unguiculata* [L.] Walp., diploid, 2*n* = 22) is a major crop grown worldwide for food and nutritional security (Lonardi et al., 2019). It is well adapted to hot, semi-arid environments, and is highly drought and heat tolerant (Hall et al., 1997). Annual cowpea production is estimated at 7 million tons of dry grain harvested on about 14 million hectares worldwide (Singh, 2020). It is grown in over two-thirds of the developing world where it is a major source of protein for human consumption, fodder for livestock (Tarawali et al., 1997), and provides ecosystem services as a cover crop to enhance soil fertility and suppresses weeds (Martins et al., 2003; Rodrigues et al., 2013). Well-fed livestock provide meat, milk, traction, and manure that contributes towards the sustainability of farming systems (Kristjanson et al., 2001). More importantly, cowpea forms a symbiotic association with root nodulating bacteria and fixes nitrogen directly to the soil (Martins et al., 2003). This biological nitrogen fixation improves crop growth and grain production without increasing production costs associated with application of nitrogen fertilizers. Crop rotation including cowpea also helps to decrease instances of *Striga hermonthica*, a parasitic weed of cereals (Berner et al., 1996).

Plant breeders exploit germplasm diversity to generate phenotypic variation for traits under selection, primarily for those influenced by climate variability (Brummer et al., 2011). Therefore, genetic and phenotypic characterization of germplasm collections is critical to warrant the development of resilient varieties that will sustain production under future scenarios of climate change. Previous cowpea genetic diversity study using a GoldenGate genotyping assay consisting of 1,536 single nucleotide polymorphisms (SNP)s on 442 cowpea landraces revealed the presence of two major gene pools in cultivated cowpea in Africa (Huynh et al., 2013). A diverse set of 768 cultivated cowpea genotypes from 58 countries were also studied using SNP markers from genotyping by sequencing (GBS) that divided the population into 3 gene pools (America, Africa, and Central West Asia) (Xiong et al., 2016). Lastly, a set of 368 cultivated cowpeas genotyped with 51,128 SNPs revealed six major subpoulations (Muñoz-Amatriaín et al., 2021). Large collections of diverse cowpea accessions are conserved in the International Institute of Tropical Agriculture (IITA) (~15,000 accessions), United States Department of Agriculture – Genetic Resources Information Network (USDA-GRIN) (7,737 accessions), and University of California, Riverside, USA (~6,000 accessions). The large number of conserved accessions in gene bank precludes their direct utilization in a breeding program owing to resource limitations in characterizing the whole collection. Therefore, a mini-core collection consisting 298 lines from the IITA collection were genotyped based on genotyping by sequencing (GBS) using 2,276 SNP markers in order to make the characterization and utilization of the germplasm more practical (Fatokun et al., 2018). Similarly, another mini-core collection, the ‘UCR Minicore’, consisting of 368 accessions that included landraces and breeding materials from 50 countries was also developed (Muñoz-amatriaín et al.) and genotyped using a publicly available Cowpea iSelect Consortium Array (Muñoz-Amatriaín et al., 2017). This array consists of 51,128 assays developed from sequencing 36 diverse accessions and was released to facilitate easy high-throughput genotyping in cowpea (Muñoz-Amatriaín et al., 2017). While progress has been made through conventional breeding in cowpea, the availability of these new molecular genetic tools enables application of modern breeding strategies for cowpea improvement (Gupta et al., 2014).

Flowering time is a key player in plant adaptation and is an important phenological trait to breed for because agronomic traits such as plant growth, plant height, and grain quality depend on the timing of flowering (Durand et al., 2012; González et al., 2016). Early flowering plants could mature earlier and help plants to avoid terminal drought stress (Kumar and Abbo, 2001). Crop legumes show large variation in flowering time, which has aided their improvement using selection and breeding (Weller and Ortega, 2015). High heritability estimates for days to flowering are reported in legumes, ranging from broad sense heritability on an entry-mean basis of 0.77 to 0.95 in soybean (Zhang et al., 2015; Mao et al., 2017), 0.38-0.75 in alfalfa (Adhikari et al., 2019), and narrow-sense heritability of 0.63 - 0.86 in cowpea (Ishiyaku et al., 2005). In many species, flowering is induced in response to day length. Different flowering responses are categorized as short-day, long-day, intermediate-day, or day-neutral based on the day length requirement to induce flowering (Bastow and Dean, 2002). Most cowpea genotypes are short-day, in which flowering is favored by day lengths shorter than the corresponding nights, while some genotypes are insensitive to a wide range of photoperiods (Summerfield and Roberts, 1985). Warmer temperatures can hasten the appearance of flowers in both daylength-sensitive and insensitive genotypes (Summerfield and Roberts, 1985). The critical photoperiod for cowpea at 27°C was reported to be between 12 and 13 h day^-1^ (Craufurd et al., 1996).

Owing to the importance of flowering time in cowpea, studies in the past have focused on identifying quantitative trait locus (QTL) using SNP and simple sequence repeat (SSR) markers in recombinant inbred lines (RIL). Five QTLs related to time of flower opening and three QTLs related to days to flower were identified in a RIL population of 524B x 219-01 using SSR markers (Andargie et al., 2013). SNP and SSR markers were utilized in another RIL population of ZN016 x ZJ282 to identify QTLs for days to first flowering, nodes to first flower, leaf senescence, and pod number per plant (Xu et al., 2013). One major QTL and few minor QTLs were found to dominate each of the four traits with three to four QTLs controlling individual traits. Other studies aimed at deciphering the genetics of flowering time in cowpea have proposed one-gene (Sène, 1967) and seven-gene (Ishiyaku et al., 2005) models to control flowering. Recent advances in genomic technologies has enabled a better understanding of the genetic basis of variation using GWAS, as it can be used for identification and high resolution mapping of useful genetic variability from germplasm sets that have resulted from many rounds of historical recombination (Yu and Buckler, 2006). GWAS studies have been reported in cowpea for pod length (Xu et al., 2017), root architecture (Burridge et al., 2017), black seed coat color (Herniter et al., 2018), seed weight, length, width, and density (Lo et al., 2019), and plant productivity traits and flowering time (Muñoz-Amatriaín et al., 2021). The study of Muñoz-Amatriaín et al. (2021) evaluated flowering time in five different environments in Nigeria and California, most of which were short-day environments.

Existing genetic diversity of cowpea needs to be assessed in order to strengthen breeding programs for developing high yielding dual-purpose cultivars with good grain and fodder yields. In this study, we phenotyped the UCR Minicore in Ft. Collins, CO and Citra, FL and performed GWAS for days to flowering; and identified candidate genes related to flowering time in cowpea.

## 2 Materials and methods

### 2.1 Germplasm, Site Description, and Experimental Design

A total of 367 accessions from the cowpea UCR Minicore (Muñoz-Amatriaín et al., 2021) were planted in Ft. Collins, Colorado (40.6553°N, −104.9966°W) on June 17, 2019. This collection includes landraces and breeding materials from 50 countries. Seeds from each accession were planted in 6.4 m rows with 0.9 m alley and 50 seeds per plot. The experiment was set up as row/column design with one replication and augmented representation of two control lines (CB5 and CB46). Plots were irrigated at the rate of 25.4 mm every week. The experiment was repeated in 2020 when the plots were planted on June 5, 2020.

A total of 292 cowpea accessions from the cowpea mini-core collection that had mature pods in October 2017 were selected from a UC-Riverside field location and planted in the field at the Plant Science Research and Experimental Unit (PSREU), Citra, FL (29.4119°N; 82.1098°W) on September 7, 2018 (Dareus et al., 2021). The soil was a Chipley sand (thermic, coated Aquic Quartzipsamments) with a pH of 6.9 and characterized by high P_2_O_5_ content, and low K_2_O, S, and Mg content. Seeds from each accession were planted in single row of 10 plants per plot, and the experiment was set up as a row/column design with two replications and augmented representation of ten control lines. Each experimental unit (3 m x 0.6 m) consisted of ten plants manually seeded and spaced at 0.3 m within row and 0.6 m between row spacing (Dareus et al., 2021).

### 2.2 Phenotypic trait and analyses

Days to flowering in Colorado was taken as the number of days from seeding to first time 50% of the plants of a given accession flowered. In Florida, days to flowering was monitored every two days, and days to first flowering was counted as the number of days from planting to the day when at least 10% of the plants in the experimental unit exhibited flowers. Descriptive analysis, and analysis of variance (ANOVA) were conducted in the R statistical package (R Development Core Team, 2020). Variance components were estimated using mixed linear models in ASReml-R v.4 (Butler et al., 2017). Best linear unbiased estimate (BLUE) and best linear unbiased prediction (BLUP) for each trait was extracted for every accession using ASREML-R (Butler et al., 2017). Broad-sense heritability (*H^2^*) was calculated using variance components by the formula H^2^=VG/(VG+VE), where VG represents genetic variance and VE represents the residual variance.

### 2.3 SNP genotyping

SNP genotyping is previously described (Muñoz-Amatriaín et al., 2017). Briefly, total genomic DNA from single plants was extracted from dried leaves using Plant DNeasy (Qiagen, Germany) and genotyped using the Cowpea iSelect Consortium Array that contained 51,128 SNPs. SNPs were called using GenomeStudio software V.2011.1 (Illumina, Inc. San Diego, CA) and the physical positions of the SNPs were determined by using the IT97K-499-35 reference genome v1.0 (Lonardi et al., 2019).

### 2.4 Genome-wide association study

Marker trait association (MTA) using all SNP markers were evaluated based on the BLUE values for days to flower. A minor allele frequency (MAF) threshold of 5% was used to remove rare variants and avoid false-positive associations. Multiple algorithms were applied for GWAS. For all SNP loci and phenotypic data, we applied the generalized linear model (GLM) and mixed linear model (MLM) implemented in GAPIT (Lipka et al., 2012). Further, GWAS was conducted using Fixed and random model Circulating Probability Unification (FarmCPU) algorithm that takes into account the confounding problem between covariates and test marker by using both Fixed Effect Model (FEM) and a Random Effect Model (REM) (Liu et al., 2016). GWAS was also conducted using BLINK that uses Bayesian Information Content (BIC) in a fixed effect model and replaces the bin approach used in FarmCPU with linkage disequilibrium (Huang et al., 2018b). Six principal components from GAPIT were used as covariates to control for population structure and manhattan plots were drawn using package *qqman* (Turner, 2014) in R statistical package (R Development Core Team, 2020).

### 2.5 Candidate gene identification

For candidate gene identification, the reference genome of cowpea IT97K-499-35 v1.0 (Lonardi et al., 2019) and the corresponding annotation (Vunguiculata_469_v1.1.annotation_info.csv) and gff file (vigna_genesv1_1_gff.csv) were used. A region of 270 kb above and below the significant SNPs was further evaluated and gene models were extracted to identify candidate genes. Orthologs of these genes on *Arabidopsis* were identified and functionally characterized using TAIR database (www.arabidopsis.org) and their molecular functions were elucidated. Gene models whose Gene Ontology (GO) function was related to flowering were selected as candidate genes and their function was searched in the literature.

### 3 Results

#### 3.1 Phenotypic Analysis

There was a significant variation in days to flower in all the datasets evaluated (Table 1, Figure 1). In Colorado in 2019, the average days to flowering was 75 days and in 2020 it was 72 days. Days to flowering was much earlier in Florida. In Florida in 2018, the average days to flowering was 41 days with a range of 32-69 days. Range of flowering was also shorter in Florida as compared to Colorado. Broad-sense heritability for flowering time ranged from 0.72-0.95 for the three studies. Pearson’s correlation between the BLUEs for the three datasets were positive (0.44-0.81) (*p* < 0.05) showing that early flowering lines in Florida also flowered early in Colorado in both years.

**Table 1.** Estimates of genotypic (s^2^g) and residual (s^2^e) variance components, broad-sense heritability (*H^2^*), standard error (SE) of the *H^2^*, number of accessions planted, mean, and range for days to flowering in the three studies.

| Location | Year | Accessions<br>Evaluated | Mean | Range | $H^2 \pm \text{SE}$ | $s^2_g$ | $s^2_e$ |
| --- | --- | --- | --- | --- | --- | --- | --- |
| Colorado | 2019 | 367 | 75 | 56-100 | $0.95 \pm 0.04$ | 102.62*** | 4.07 |
| Colorado | 2020 | 367 | 72 | 50-88 | $0.80 \pm 0.12$ | 47.62* | 11.01 |
| Florida | 2018 | 292 | 41 | 32-69 | $0.72 \pm 0.06$ | 11.01*** | 2.93 |

‘\*\*\*’ denotes significance at $p < 0.001$ and ‘\*’ denotes significance at $p < 0.05$ for the Likelihood Ratio Tests (LRT)

**Figure 1:**
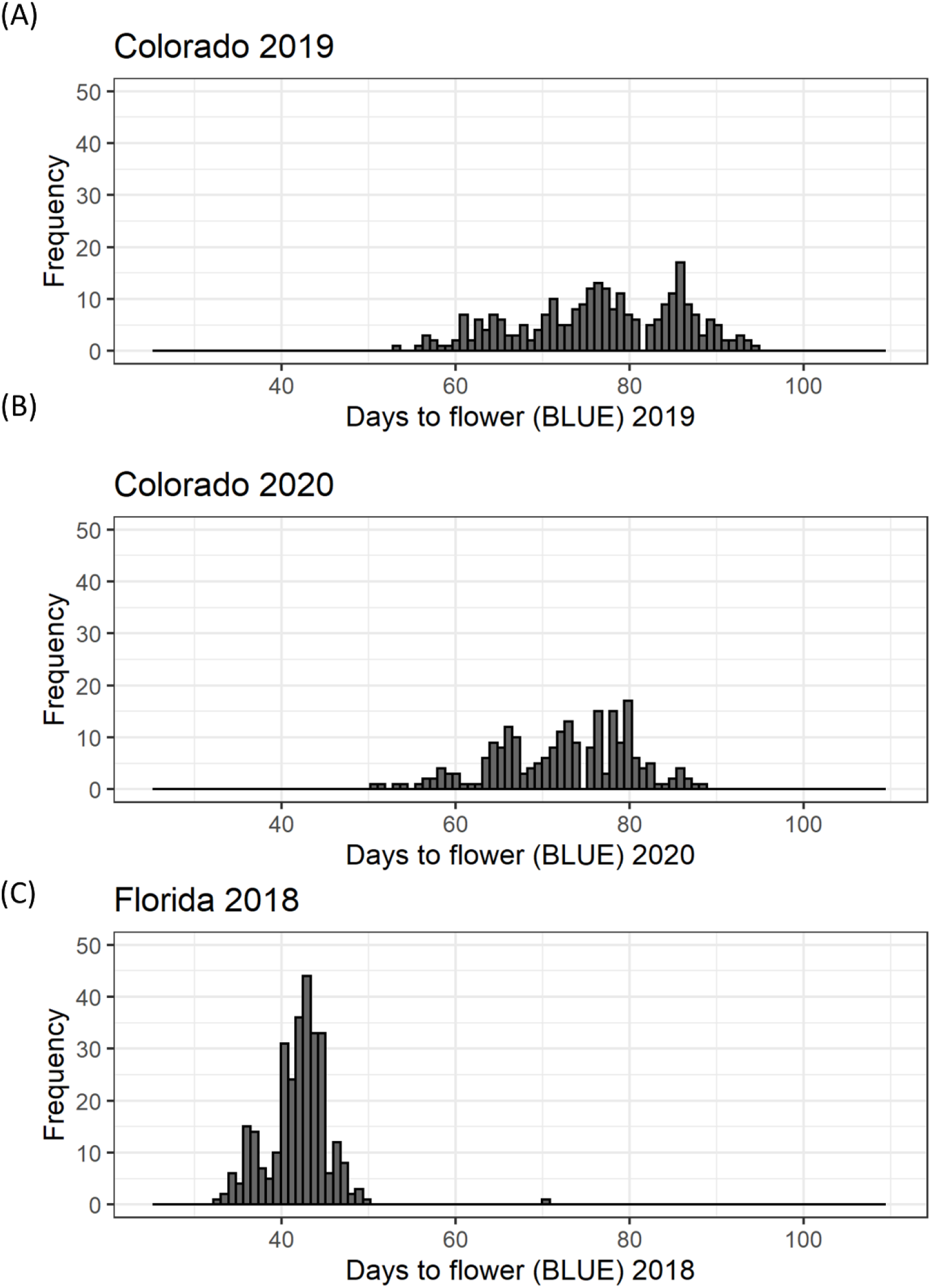
Histogram of Days to flower for: (A) 367 accessions of the cowpea mini-core collection planted in 2019 in Colorado; (B) 367 accessions of the cowpea mini-core collection planted in 2020 in Colorado; and (C) 292 accessions of the cowpea mini-core collection planted in 2018 in Florida.

#### 3.2 Weather data

Daily maximum and minimum temperatures were lower in Colorado than in Florida (Supplementary Figure S1, Supplementary Figure S2, and Supplementary Figure S3). Minimum day length during the experimental period in Colorado was 12.05 hours in 2019 and 12.85 hours in 2020 while that was 10.25 hours in Florida in 2018. Daylength was slowly decreasing from planting to flowering in all the trials. In Colorado, the minimum daylength when the first plots had 50% flowering was 13.9 hours with an average temperature of 22.9°C in 2019 and 14.52 hours with average temperature of 22.8°C in 2020. Minimum daylength when the first flowering occurred in Florida in 2018 was 11.65 hours with average temperature of 26.6°C.

#### 3.3 Genome wide association studies

All SNP markers after filtering for MAF were used for GWAS. We identified 30 MTAs corresponding to 20 unique SNPs for days to flowering that explained 8-12% of phenotypic variance in the GWAS conducted using four software in the three datasets (Table 2). These significant MTAs were distributed across seven chromosomes of the cowpea genome (Figure 2, Supplementary Figure S4-S6). In chromosome Vu03, FarmCPU identified a single SNP (2_03926). Multiple MTAs were identified on chromosome Vu04. FarmCPU, BLINK, and GLM identified the same significant SNP in chromosome Vu04 (2_55402), while both BLINK and GLM identified SNP 2_06977. GLM, FarmCPU, and BLINK further identified 7, 1, and 2 additional unique MTAs respectively, on chromosome Vu04 (Table 2). FarmCPU identified two unique MTAs (2_42453 and 2_43970) on chromosome Vu07. In chromosome Vu08, FarmCPU identified the same SNP (1_0362) in two studies (Colorado 2019 and Colorado 2020). FarmCPU further identified two unique MTAs in chromosome Vu09 and one unique MTA each in chromosome Vu10 and chromosome Vu11. BLINK identified one unique MTA in chromosome Vu10 (2_54017). MLM did not identify any significant MTAs in the three GWAS studies. Seven unique markers were reliable as they were identified by multiple algorithms or identified in more than 1 GWAS study (Table 2). In Colorado in 2019, early flowering alleles decreased flowering time by 5.50-6.93% corresponding to an average number of 4-6 days (Figure 3). Similarly, in Colorado in 2020, early flowering alleles decreased flowering time by 5.06-6.74% corresponding to an average number of days to 4-5 days. In Florida in 2018, early flowering alleles decreased flowering time by 6.32% corresponding to a decrease in flowering by 3 days.

**Table 2.** Significant SNPs related to days to flowering identified by multiple algorithms in genome wide association studies in the three studies along with their *p* value, minor allele frequency (MAF), effect, percentage of variance explained (PVE(%)) as reported by each software, and -log_10_(*p*).

| SNP | Chr. | Position | $p\_value$ | MAF | Efect | Location | Year | Software | PVE (%) | $-\log_{10}(p)$ |
| --- | --- | --- | --- | --- | --- | --- | --- | --- | --- | --- |
| 2_03926 | Vu03 | 50079486 | 7.10E-07 | 0.22 | -2.36 | Colorado | 2019 | FarmCPU | NA | 6.15 |
| 2_33309 | Vu04 | 9183563 | 2.34E-07 | 0.11 | -5.00 | Colorado | 2019 | GLM | 9.20 | 6.63 |
| 2_06977 | Vu04 | 9211195 | 2.68E-14 | 0.11 | NA | Colorado | 2019 | BLINK | NA | 13.57 |
| 2_06977 | Vu04 | 9211195 | 3.28E-09 | 0.11 | -5.98 | Colorado | 2019 | GLM | 12.25 | 8.48 |
| 2_06977 | Vu04 | 9211195 | 5.37E-12 | 0.09 | NA | Colorado | 2020 | BLINK | NA | 11.27 |
| 2_06977 | Vu04 | 9211195 | 8.24E-08 | 0.09 | -4.94 | Colorado | 2020 | GLM | 11.32 | 7.08 |
| 2_52931 | Vu04 | 9224808 | 4.13E-08 | 0.09 | -5.62 | Colorado | 2019 | GLM | 10.42 | 7.38 |
| 2_52931 | Vu04 | 9224808 | 1.03E-07 | 0.09 | -4.89 | Colorado | 2020 | GLM | 11.13 | 6.99 |
| 2_05817 | Vu04 | 9263427 | 3.13E-08 | 0.10 | 5.61 | Colorado | 2019 | GLM | 10.62 | 7.50 |
| 2_05817 | Vu04 | 9263427 | 1.53E-07 | 0.09 | 4.84 | Colorado | 2020 | GLM | 10.81 | 6.81 |
| 2_04510 | Vu04 | 10681090 | 5.14E-08 | 0.09 | 5.69 | Colorado | 2019 | GLM | 10.27 | 7.29 |
| 2_04510 | Vu04 | 10681090 | 5.64E-07 | 0.08 | 4.71 | Colorado | 2020 | GLM | 9.77 | 6.25 |
| 2_38629 | Vu04 | 10776312 | 1.09E-06 | 0.11 | -4.54 | Colorado | 2019 | GLM | 8.12 | 5.96 |
| 2_38629 | Vu04 | 10776312 | 1.04E-06 | 0.10 | -4.14 | Colorado | 2020 | GLM | 9.28 | 5.98 |
| 2_46442 | Vu04 | 20308708 | 1.01E-06 | 0.38 | 3.17 | Colorado | 2019 | GLM | 8.18 | 6.00 |
| 2_55402 | Vu04 | 27032485 | 3.10E-11 | 0.25 | -1.16 | Florida | 2018 | FarmCPU | NA | 10.51 |
| 2_55402 | Vu04 | 27032485 | 3.65E-13 | 0.25 | NA | Florida | 2018 | BLINK | NA | 12.44 |
| 2_55402 | Vu04 | 27032485 | 2.40E-08 | 0.25 | -1.41 | Florida | 2018 | GLM | 10.11 | 7.62 |
| 2_52369 | Vu04 | 27310629 | 2.93E-07 | 0.33 | NA | Colorado | 2020 | BLINK | NA | 6.53 |
| 2_22451 | Vu04 | 27793336 | 4.77E-07 | 0.18 | 1.45 | Florida | 2018 | GLM | 8.15 | 6.32 |
| 2_27454 | Vu04 | 32104992 | 4.58E-07 | 0.29 | 2.31 | Colorado | 2019 | FarmCPU | NA | 6.34 |
| 2_42453 | Vu07 | 28483321 | 3.75E-07 | 0.36 | -2.26 | Colorado | 2019 | FarmCPU | NA | 6.43 |
| 2_43970 | Vu07 | 38052504 | 6.80E-07 | 0.14 | -2.76 | Colorado | 2019 | FarmCPU | NA | 6.17 |
| 1_0362 | Vu08 | 29639172 | 2.61E-07 | 0.45 | 1.83 | Colorado | 2019 | FarmCPU | NA | 6.58 |
| 1_0362 | Vu08 | 29639172 | 7.07E-07 | 0.45 | 1.47 | Colorado | 2020 | FarmCPU | NA | 6.15 |
| 2_39424 | Vu09 | 419806 | 1.62E-08 | 0.45 | -0.79 | Florida | 2018 | FarmCPU | NA | 7.79 |
| 2_04844 | Vu09 | 6752951 | 9.22E-11 | 0.27 | 2.97 | Colorado | 2020 | FarmCPU | NA | 10.04 |
| 2_54017 | Vu10 | 7807120 | 2.99E-08 | 0.26 | NA | Colorado | 2019 | BLINK | NA | 7.52 |
| 2_42049 | Vu10 | 13068722 | 3.92E-08 | 0.12 | 4.76 | Colorado | 2020 | FarmCPU | NA | 7.41 |
| 2_03469 | Vu11 | 30442337 | 4.48E-07 | 0.18 | -1.96 | Colorado | 2020 | FarmCPU | NA | 6.35 |

**Figure 2:**
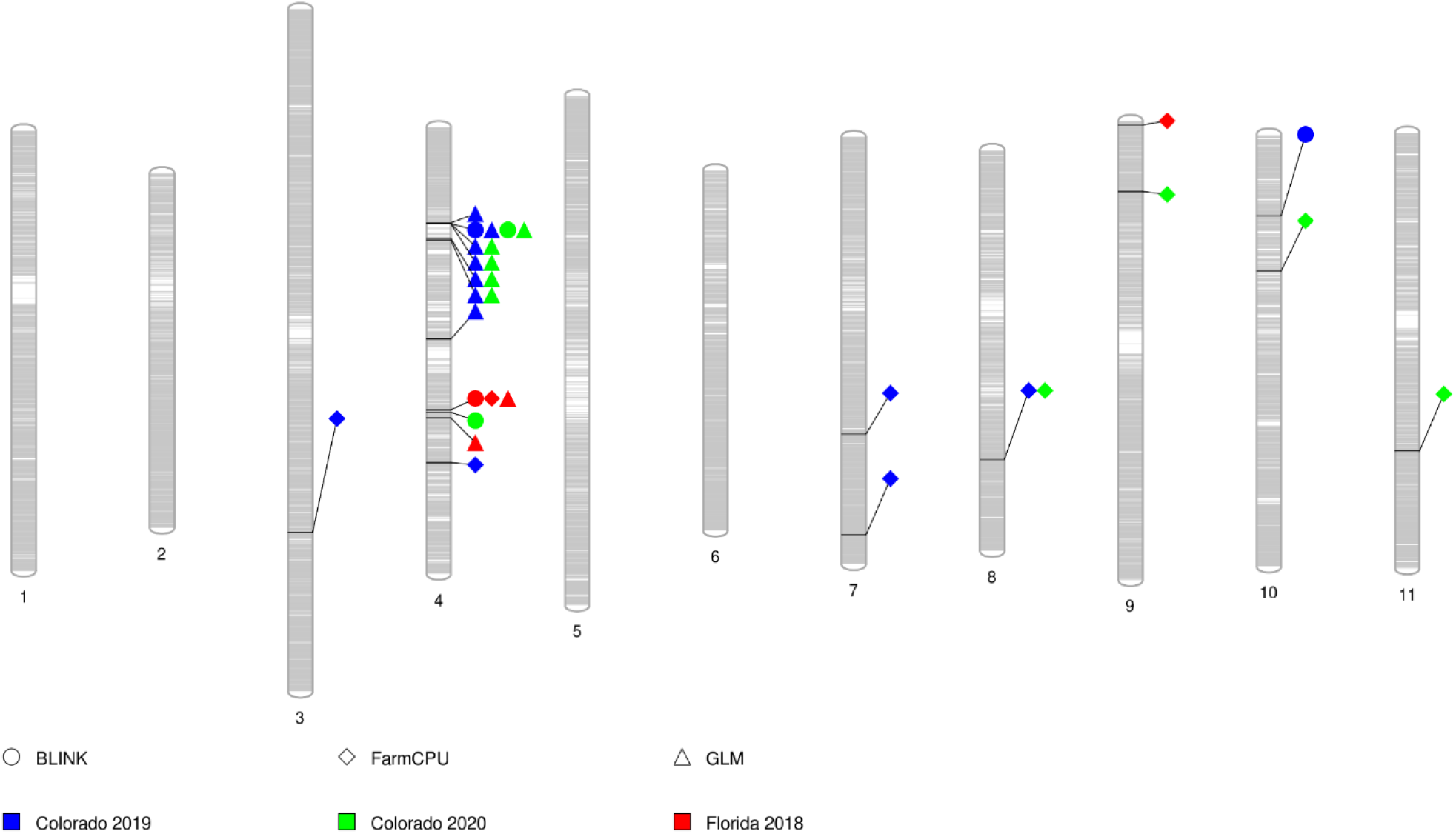
PhenoGram showing significant marker-trait associations for flowering time on each chromosome. The grey bars within each chromosome show the locus of SNPs in the chromosome. Each shape represents a significant SNP identified by the three algorithms (circle = BLINK, diamond = FarmCPU, and triangle = GLM). The color within each shape represents SNPs identified in the different studies (red = Florida 2018, blue = Colorado 2019, and green = Colorado 2020).

**Figure 3.**
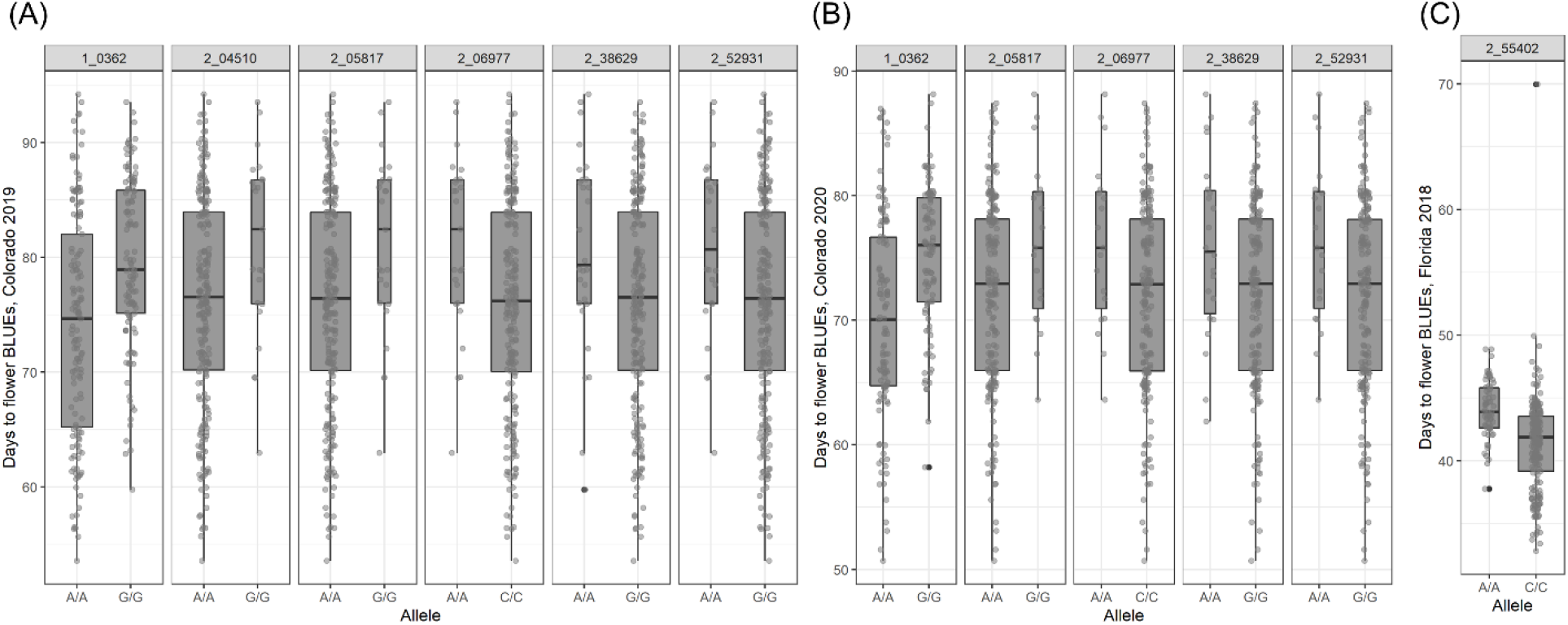
Boxplot of days to flower as affected by the alleles present on the population (A) 367 accessions of the cowpea mini-core collection planted in 2019 in Colorado; (B) 367 accessions of the cowpea mini-core collection planted in 2020 in Colorado; and (C) 292 accessions of the cowpea mini-core collection planted in 2018 in Florida.

#### 3.4 Candidate gene identification

The linkage region of the 20 significant SNPs (SNP ± 270 kb) harbored a total of 483 unique gene models on the cowpea genome. Functional annotation of these gene models using the *Arabidopsis* gene network identified a total of 12 genes that were related to flowering (Table 3). These genes included important genes like FLOWERING LOCUS T (*FT*), GIGANTEA (*GI*), Cryptochrome-2 (*CRY2*), LIGHT-DEPENDENT SHORT HYPOCOTYLS 3 (*LSH3*), REBELOTE (*RBL*) that are known to control flowering time in *Arabidopsis* and other species (El-Assal et al., 2001; Teper-Bamnolker and Samach, 2005; Prunet et al., 2008; Takeda et al., 2011; Park et al., 2020). These candidate genes were located in chromosomes Vu04, Vu07, Vu08, and Vu09. In chromosome Vu04, the peak signal at locus 2_46442 was associated with *RBL* gene, 2_55402 was associated with *FT* gene, and 2_27454 was associated with *GI* gene. In chromosome Vu07, the peak signal at locus 2_42453 was associated with two genes *CRY2* and *LSH3*. In chromosome Vu08, locus 1_0362 was tied to three genes: *UGT87A2, BBX32* and Snf1 kinase interactor-like protein. Finally, in chromosome Vu09, locus 2_39424 was associated with *NGA1, DCL1*, and *LIF2* while locus 2_04844 was associated with *HTA9*.

**Table 3.** Genes related to flowering time that are within ±270kb of the significant SNPs. Locus name is the name of the gene in the cowpea reference genome with start and end for each gene in the chromosome, name of the associated SNP, position of the SNP, associated gene name, and locus ID of the associated gene in *Arabidopsis*.

| Locus Name | Chr | Start | End | SNP | Position | Gene Name | *Arabidopsis* locus ID |
| --- | --- | --- | --- | --- | --- | --- | --- |
| Vigun04g096400 | Vu04 | 20505963 | 20516534 | 2_46442 | 20308708 | *RBL* | AT3G55510 |
| Vigun04g109500 | Vu04 | 27156677 | 27161340 | 2_55402 | 27032485 | *FT* | AT1G65480 |
| Vigun04g126700 | Vu04 | 32034709 | 32050305 | 2_27454 | 32104992 | *GI* | AT1G22770 |
| Vigun07g171300 | Vu07 | 28639274 | 28644217 | 2_42453 | 28483321 | *CRY2* | AT1G04400 |
| Vigun07g171900 | Vu07 | 28713837 | 28716092 | 2_42453 | 28483321 | *LSH3* | AT2G31160 |
| Vigun08g124100 | Vu08 | 29426933 | 29428985 | 1_0362 | 29639172 | *UGT87A2* | AT2G30140 |
| Vigun08g127400 | Vu08 | 29776374 | 29777564 | 1_0362 | 29639172 | *BBX32* | AT3G21150 |
| Vigun08g128600 | Vu08 | 29870661 | 29873299 | 1_0362 | 29639172 | *Snf1* kinase | AT1G80940 |
| Vigun09g003600 | Vu09 | 249165 | 253046 | 2_39424 | 419806 | *NGA1* | AT2G46870 |
| Vigun09g003800 | Vu09 | 275035 | 288027 | 2_39424 | 419806 | *DCL1* | AT1G01040 |
| Vigun09g005800 | Vu09 | 426595 | 430905 | 2_39424 | 419806 | *LIF2* | AT4G00830 |
| Vigun09g063700 | Vu09 | 6692636 | 6694999 | 2_04844 | 6752951 | *HTA9* | AT1G52740 |

### 4 Discussion

This study evaluated the variation in flowering time in the cowpea UCR Minicore in two contrasting environments in Colorado and Florida. There was a wide variation in days to flower in all trials. We observed high *H^2^* estimates (0.72-0.95) for flowering time in cowpea, which is similar to the estimates reported in other species like soybean (0.77-0.95) (Zhang et al., 2015; Mao et al., 2017), and alfalfa (0.38-0.75) (Adhikari et al., 2019). High *H^2^* of flowering time shows the inherent genetic control of flowering as seen in other species. A *H^2^* of 84.5% was reported for days to flower in cowpeas (Omoigui et al., 2006) and a narrow-sense heritability (*h^2^*) of 86% was reported in a cross between photoperiod-sensitive and photoperiod-insensitive varieties with at least seven major gene pairs estimated to control time of flowering in this population (Ishiyaku et al., 2005). Since flowering time is an important trait for plant breeders, the presence of variation in flowering time for cowpea shows a large potential to manipulate its expression by breeding and selection.

Flowering time is a complex trait (Weller and Ortega, 2015) and is generally regulated by genetic networks composed of four main converging pathways: autonomous, gibberellin, photoperiod, and vernalization (Roux et al., 2006). These pathways integrate physiological and environmental cues to activate the transition from vegetative to reproductive stages at an optimum time (Brock et al., 2009). In *Arabidopsis*, induced mutations revealed the existence of up to 80 loci that affected flowering time (Levy and Deant, 1998). In cowpea, previous studies aimed at elucidating the genetics of flowering time have mostly focused on QTL analysis. Three QTLs related to days to flower and five QTLs related to time of flower opening were identified using 202 SSR markers in a mapping population of 159 F7 lines obtained by crossing a short duration variety (524B) to a long duration variety (219-01) (Andargie et al., 2013). The linkage groups in this study were not named based on the reference genome (Lonardi et al., 2019), therefore, these QTLs could not be directly compared with our results. SNP and SSR markers were utilized in another RIL population of ZN016 × ZJ282 to identify QTLs for days to first flowering, nodes to first flower, leaf senescence, and pod number per plant (Xu et al., 2013). One major QTL and few minor QTLs were found to dominate each of the four traits with three to four QTLs controlling individual traits. Similarly, two QTLs on chromosome Vu05 and chromosome Vu09 with peak SNPs at 2_05332 (854,745 bp) and 2_03945 (5,449,874 bp) respectively, were identified for days to flowering using 215 F8 RILs derived from a cross between cultivated (IT99K-573-1-1) and wild (TVNu-1158) cowpea accession (Lo et al., 2018). Studies on the cowpea multi-parent advanced generation intercross (MAGIC) population have identified flowering time loci with up to 25% phenotypic variability explained (PVE) and additive effect size of 7 days under long-days but not under short-days (Olatoye et al., 2019). Drought tolerance index for flowering time in this population identified significant SNPs (2_06470, 2_52919, 2_06137, and 1_0946) on chromosome Vu03 that were 12Mb downstream of the significant SNP identified in our study (2_03926) (Ravelombola et al., 2021). Researchers have proposed one-gene (Sène, 1967) and seven-gene (Ishiyaku et al., 2005) models to control flowering in cowpea and suggest that distinct and common genetic regulators control flowering time adaptation to both long- and short-day photoperiod in cowpea (Olatoye et al., 2019). Few GWAS have been reported in cowpea for pod length (Xu et al., 2017), root architecture (Burridge et al., 2017), black seed coat color (Herniter et al., 2018), and seed weight, length, width, and density (Lo et al., 2019). The availability of the reference genome of cowpea and the Cowpea iSelect Consortium Array have opened up new avenues in cowpea genetic analysis (Lonardi et al., 2019). The Cowpea iSelect Consortium Array with 51,128 SNPs is an excellent tool to identify marker trait associations and population genetic studies in cowpea (Huang et al., 2018a).

For GWAS, we used four algorithms implemented in GAPIT (Lipka et al., 2012), namely, GLM, MLM, FarmCPU (Liu et al., 2016), and BLINK (Huang et al., 2018b). In a GLM, false positives are eliminated by fitting population structure as covariate (Price et al., 2006) and in MLM, population structure and genetic effect of each individual is fitted as covariates (Yu et al., 2006). FarmCPU performs marker tests with associated markers as covariates in a fixed effect model (Liu et al., 2016) and assumes that quantitative trait nucleotides (QTN) underlying the trait are distributed equally across the genome. Optimization on the associated covariate markers is done separately in a random effect model. On the other hand, BLINK eliminates the requirement of equal distribution of QTNs by taking linkage disequilibrium into consideration (Huang et al., 2018b). It also replaces the Restricted Maximum Likelihood (REML) in the mixed linear model in FarmCPU with Bayesian Information Content (BIC) in a fixed effect model to boost computing speed. These algorithms identified multiple MTAs for flowering time that were distributed in seven chromosomes in the cowpea genome.

Seven significant SNPs identified in our study harbored important flowering time related genes. On chromosome Vu04, *RBL* gene was 197 kb upstream of the significant SNP (2_46442) and this gene redundantly influences floral meristem termination (Prunet et al., 2008). *FT* was located 124 kb downstream of the most significant SNP (2_55402). *FT*, together with LEAFY (*LFY*), integrates environmental signaling for induction of flowering (Moraes et al., 2019). *Arabidopsis FT* is a member of a six-gene family that includes another important flowering-related gene, TERMINAL FLOWER1 (*TFL1*) that delays transition to flowering and has been identified in legumes like pea, *Medicago*, and lotus (Hecht et al., 2005). *FT* is expressed in leaves and is induced by long-day treatment in *Arabidopsis* (Teper-Bamnolker and Samach, 2005). Additionally, in chromosome Vu04, *GI* was located 70 kb downstream of the most significant SNP (2_27454). GI-mediated integration of photoperiodic and temperature information shapes thermo-morphogenic adaptation responses in plants that optimizes plant growth and fitness in warm climates (Park et al., 2020). A total of 11 SNPs significantly associated with flowering time were identified in chromosome Vu04 showing that this chromosome is very important in cowpea for adaptation and selection for flowering. On chromosome Vu07, SNP 2_42453 harbored multiple genes. *CRY2* was located 155 kb downstream of the SNP while *LSH3* was located 230 kb downstream of the SNP. *CRY2* is a blue light receptor that mediates blue-light regulated cotyledon expansion and is involved in the flowering response to photoperiod in *Arabidopsis* (El-Assal et al., 2001). It is also a positive regulator of the flowering-time gene *CONSTANS* (Guo et al., 1998). *LSH3*, also known as *ORGAN BOUNDARY 1* encodes ALOG family proteins and is expressed at the boundary of shoot apical meristem and lateral organs (Takeda et al., 2011). Constitutive expression of *LSH3* and *LSH4* generates chimeric floral organs.

In chromosome Vu08, SNP 1_0362 harbored 3 genes: *UGT87A2* located 212 kb downstream, *BBX32* located 137 kb upstream, and Snf1 kinase interactor-like protein located 231 kb downstream of the SNP. *UGT87A2* promotes early flowering and is an important player in the autonomous pathway (Wang et al., 2012) while *BBX32* is regulated by circadian clock and regulates flowering and hypocotyl growth (Tripathi et al., 2017). In chromosome Vu09, SNP 2_39424 harbored three genes: *NGA1* located 170 kb downstream, *DCL1* located 144 kb downstream, and *LIF2* located 7 kb upstream of the SNP. *NGA* directs development of apical tissues in gynoecium (Ballester et al., 2015), *DCL1* promotes flowering by repressing *FLOWERING LOCUS C* (Schmitz et al., 2007), and *LIF2* regulates flower development and maintains ovary determinacy in short day conditions (Latrasse et al., 2011). Additionally, in chromosome Vu09, *HTA9* was located 60kb downstream of the SNP 2_04844 and this gene mediates the thermo-sensory flowering response in *Arabidopsis* (Jarillo and Piñeiro, 2015). Identification of multiple significant SNPs and genes related to flowering time in the cowpea genome suggests their important role in controlling flowering time in cowpeas as well as the complex nature of flowering time trait. These genes should be the primary targets for modifications while breeding cowpea and further detailed studies of these candidate genes will help to decipher the overall mechanism of flowering in cowpea.

MTAs in our study could not be directly compared to previous QTL studies (Andargie et al., 2013; Xu et al., 2013) because of the absence of common markers. In a previous QTL study, two significant QTLs for days to flowering were detected, one each on chromosome 5 and chromosome 9 that harbored phytochrome E and transcription factor TCP 18 that are involved in flowering time (Lo et al., 2018). Similarly, another QTL report identified three QTLs related to days to flowering, one each on LG1, LG2, and LG7 (Andargie et al., 2013). Our GWAS results detected significant reliable SNPs on chromosome Vu04 and Vu08. A recent study that utilized the SNP array in the cowpea UCR Minicore identified the same SNP (2_06977) on chromosome 4 under long days in California (Muñoz-Amatriaín et al., 2021). In our analysis, this SNP was identified by multiple algorithms in two different datasets and is most likely an important region of interest for flowering time. Interestingly, another study that utilized the SNP array identified two QTLs for flowering time in chromosomes 5 and 9 that could explain 20-79% of the phenotypic variance (Lo et al., 2018). On chromosome 9, the previously identified QTL was 1.3 Mb upstream of the SNP (2_04844) identified in this study. This suggests that these regions harbor important flowering related genes. Previous studies reported that the QTLs could explain 5-18.5% (Andargie et al., 2013), 16-30% (Xu et al., 2013), and 20-79% (Lo et al., 2018) of the phenotypic variance for days to flowering depending on the population. In our study, the variation explained by the MTAs varied from 8-12%, indicating that multiple genes might be affecting the traits and those genes have small effects. Our GWAS results in Florida were limited to accessions that flowered under the long-day conditions of Riverside (CA, USA) lines only, therefore, GWAS results from this location might miss some markers that were identified in Colorado where the whole mini-core was evaluated. Nevertheless, our study contributes with a large number of MTAs in cowpea for flowering time. Several loci identified here can be further explored for use in marker-assisted selection, genomic selection, and gene discovery.

Plant breeders develop new varieties with increased yield by improving the crop’s adaptability and stress tolerance (Brummer et al., 2011). Flowering time has been associated with adaptation and agronomic performance of traits in several crops. Early flowering plants could mature earlier and avoid drought stress. Considerable gains can be made to increase yield and stability of grain legumes in drought prone environments by shortening crop duration (Subbarao et al., 1995). This would be important in Colorado and other regions of the semi-arid High Plains, where dryland agriculture constitutes a significant proportion of the total cropland and where erratic precipitation patterns due to climate change are threatening the productivity and profitability of such system (Rosenzweig and Schipanski, 2019). Earlier flowering cowpea varieties could also help intensify dryland cropping systems in the High Plains by providing a viable alternative to the summer fallow that precedes winter wheat (Nielsen and Vigil, 2005). In the case of Florida, although the Köppen-Trewartha Climate Classification system has classified Central/North Florida as a Subtropical and Mediterranean climate, and South Florida as a Tropical climate (Belda et al., 2014), drought stress is a seasonal abiotic stressor in the state due to its sandy soil and high evaporative demand.

Early flowering can be transferred to cultivated cowpea through hybridization with early flowering accessions. Selection of early flowering cowpea that performs well in subtropical regions will undoubtedly help to increase the global production of cowpea as well as help to develop climate resilient cowpea accessions. On the other hand, extended vegetative period in late maturing varieties can provide higher biomass production which would be ideal for forage and cover crop cultivation, where the crops can be terminated before they flower and seed, thus avoiding potential invasiveness. Vegetative growth and rate of plant production have been shown to have additive and epistatic relationships with flowering time QTLs in common beans using comparative QTL mapping, suggesting pleiotropic effects between these traits (González et al., 2016). Further research is needed to identify the haplotypes that confer early or late flowering trait in cowpeas. This study established the basis for marker-assisted selection of flowering time in cowpea breeding programs. Additionally the recent availability of the reference genome (Lonardi et al., 2019), development of the cowpea UCR Minicore (Muñoz-Amatriaín et al., 2021), and future analysis of transcriptome profiles will facilitate identification and manipulation of causative loci governing flowering time across a broad range of environmental conditions.

## 5 Figure legends

Figure 1: Histogram of days to flower for the cowpea mini-core collection: (A) 367 accessions planted in 2019 in Colorado; (B) 367 accessions planted in 2020 in Colorado; and (C) 292 accessions planted in 2018 in Florida.

Figure 2: PhenoGram showing significant marker-trait associations for flowering time on each chromosome. The grey bars within each chromosome show the locus of SNPs in the chromosome. Each shape represents a significant SNP identified by the three algorithms (circle = BLINK, diamond = FarmCPU, and triangle = GLM). The color within each shape represents SNPs identified in the different studies (blue = Colorado 2019, green = Colorado 2020, and red = Florida 2018).

Figure 3. Boxplot of days to flower as affected by the alleles present on the population (A) 367 accessions of the cowpea mini-core collection planted in 2019 in Colorado; and (B) 367 accessions of the cowpea mini-core collection planted in 2020 in Colorado; and (C) 292 accessions of the cowpea mini-core collection planted in 2018 in Florida.

## 6 Table legends

Table 1. Estimates of genotypic (s^2^g) and residual (s^2^e) variance components, broad-sense heritability (*H^2^*), standard error (SE) of the *H^2^*, number of accessions planted, mean, and range for days to flowering in the three studies.

Table 2. Significant SNPs related to days to flowering identified by multiple algorithms in genome wide association studies in the three studies along with their *p* value, minor allele frequency (MAF), effect, percentage of variance explained (PVE(%)) as reported by each software, and -log_10_(*p*).

Table 3. Genes related to flowering time that are within ±270kb of the significant SNPs. Locus name is the name of the gene in the cowpea reference genome with start and end for each gene in the chromosome, name of the associated SNP, position of the SNP, associated gene name, and locus ID of the associated gene in *Arabidopsis*.

## 7 Supplementary figure legends

Supplementary Figure S1. Daily maximum (MaxT) and minimum (MinT) temperature and photoperiod (orange line) in Ft. Collins, CO during the trial in 2019.

Supplementary Figure S2. Daily maximum (MaxT) and minimum (MinT) temperature and photoperiod (orange line) in Ft. Collins, CO during the trial in 2020.

Supplementary Figure S3. Daily maximum (MaxT) and minimum (MinT) temperature and photoperiod (orange line) in Citra, FL during the trial in 2018.

Supplementary Figure S4: Manhattan plots from the GWAS analysis pertaining to 367 accessions of the cowpea mini-core collection planted in 2019 in Colorado. Left panel: Negative log_10_-transformed P values for each SNP (y axis) are plotted against the chromosomal position (y axis). The red line represents Bonferroni-corrected threshold of 0.05 for genome-wide statistically significant associations and the blue line shows suggestive associations (*p* = 1 × 10^-5^). Right panel shows the QQ plots where x-axis is expected negative log p-values and the y-axis is observed negative log p-values. GWAS results for days to flowering using (A) BLINK; (B) FarmCPU; (C) GLM; and (D) MLM.

Supplementary Figure S5: Manhattan plots from the GWAS analysis pertaining to 367 accessions of the cowpea mini-core collection planted in 2020 in Colorado. Left panel: Negative log_10_-transformed P values for each SNP (y axis) are plotted against the chromosomal position (y axis). The red line represents Bonferroni-corrected threshold of 0.05 for genome-wide statistically significant associations and the blue line shows suggestive associations (*p* = 1 × 10^-5^). Right panel shows the QQ plots where x-axis is expected negative log p-values and the y-axis is observed negative log p-values. GWAS results for days to flowering using (A) BLINK; (B) FarmCPU; (C) GLM; and (D) MLM.

Supplementary Figure S6: Manhattan plots from the GWAS analysis pertaining to 292 accessions of the cowpea mini-core collection planted in 2018 in Florida. Left panel: Negative log_10_-transformed P values for each SNP (y axis) are plotted against the chromosomal position (y axis). The red line represents Bonferroni-corrected threshold of 0.05 for genome-wide statistically significant associations and the blue line shows suggestive associations (*p* = 1 × 10^-5^). Right panel shows the QQ plots where x-axis is expected negative log p-values and the y-axis is observed negative log p-values. GWAS results for days to flowering using (A) BLINK; (B) FarmCPU; (C) GLM; and (D) MLM.

## 9 Author Contributions

ER conceived the project. ER and RD collected the phenotypic data in Florida. MM and JR collected the phenotypic data in Colorado and provided the genotypic data. DP analysed the data and wrote the manuscript. All authors reviewed the manuscript.

## 11 Acknowledgments

We would like to thank Dr. Timothy J. Close and Dr. Philip A. Roberts for providing the seeds, and Brooke Sayre-Chavez and Amanda Amsberry for their help in scoring days to flowering in Colorado. The authors thank all the Forage Breeding and Genetics Lab members and staff at the University of Florida Plant Science Research and Education Unit, Citra, FL for providing help for the field trial and data collection.

## 13 Data Availability Statement

The genetic data used in this study is available at (Muñoz-Amatriaín et al., 2021).

## 14 Contribution to the Field Statement

Plant adaptation to new climates and their yield are strongly affected by flowering time. Early flowering plants could mature earlier and avoid drought stress. This might be a good adaptation strategy to cope with impending climate change crisis, especially in regions with lower access to irrigation water. Therefore, understanding the genetic basis of flowering time is critical to advance plant breeding. Genome wide association studies for flowering time have been done in other species, however, this has not been widely reported in cowpea. Cowpea is highly heat tolerant and is an important crop to breed for new varieties that are resilient to changing climates. Cowpea is a major source of protein for human consumption as well as a quality forage for animal feed. To facilitate future plant breeding efforts, we have identified marker trait associations related to flowering time in a cowpea mini-core collection. This study contributed large number of marker trait associations in cowpea for flowering time and identified candidate genes related to flowering time in cowpea. Several loci identified here can be validated in other populations to support cowpea breeding programs with introgression of favorable alleles and marker-assisted selection, genomic selection, and gene discovery. To our knowledge, this is the first published study that has done GWAS for flowering time in cowpea using the cowpea mini-core collection.

