## Supplementary figures and images for "Genome-wide association study reveals candidate genes for flowering time in cowpea (*Vigna unguiculata* [L.] Walp)"

### Supplemental Figures

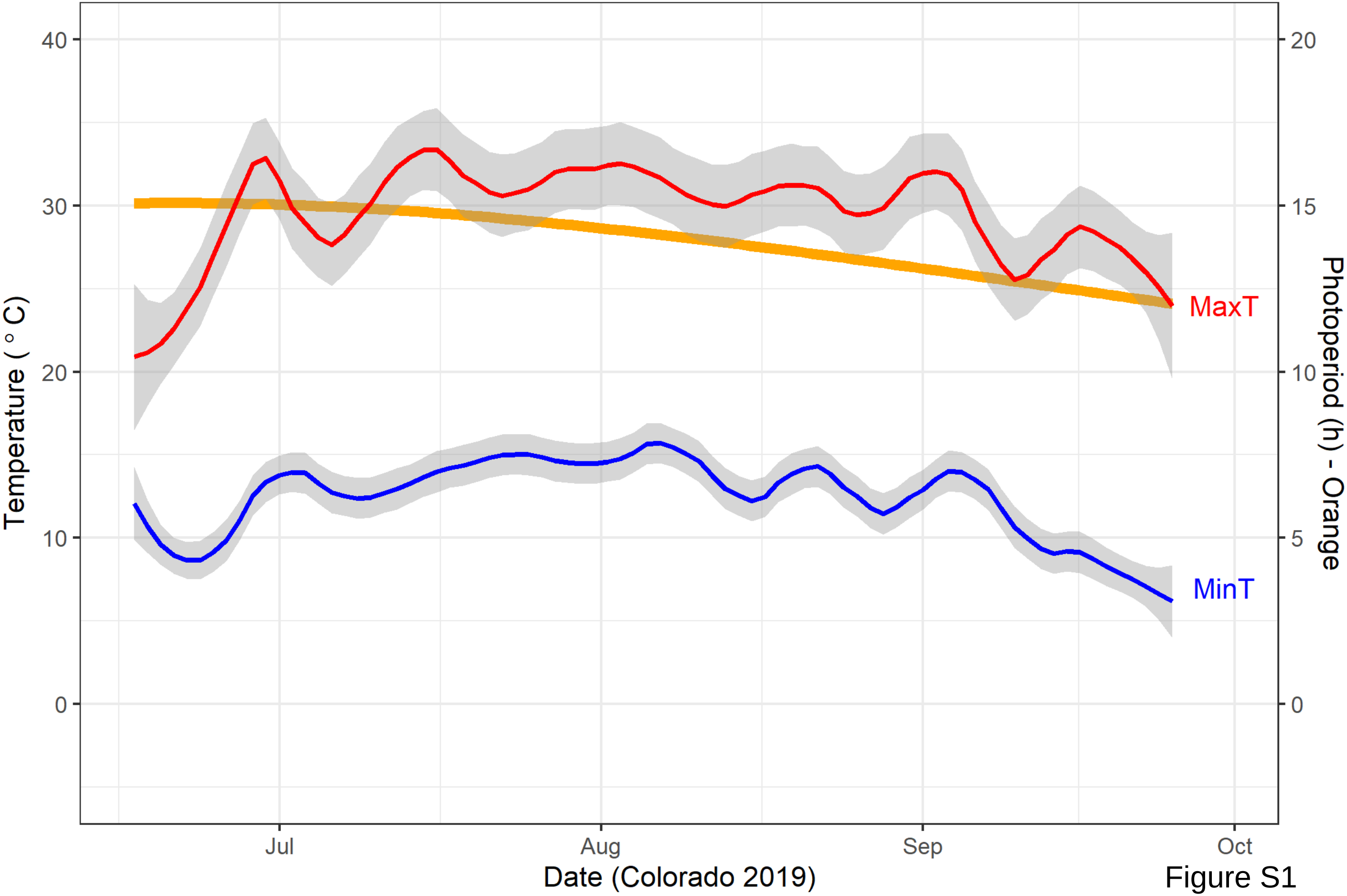

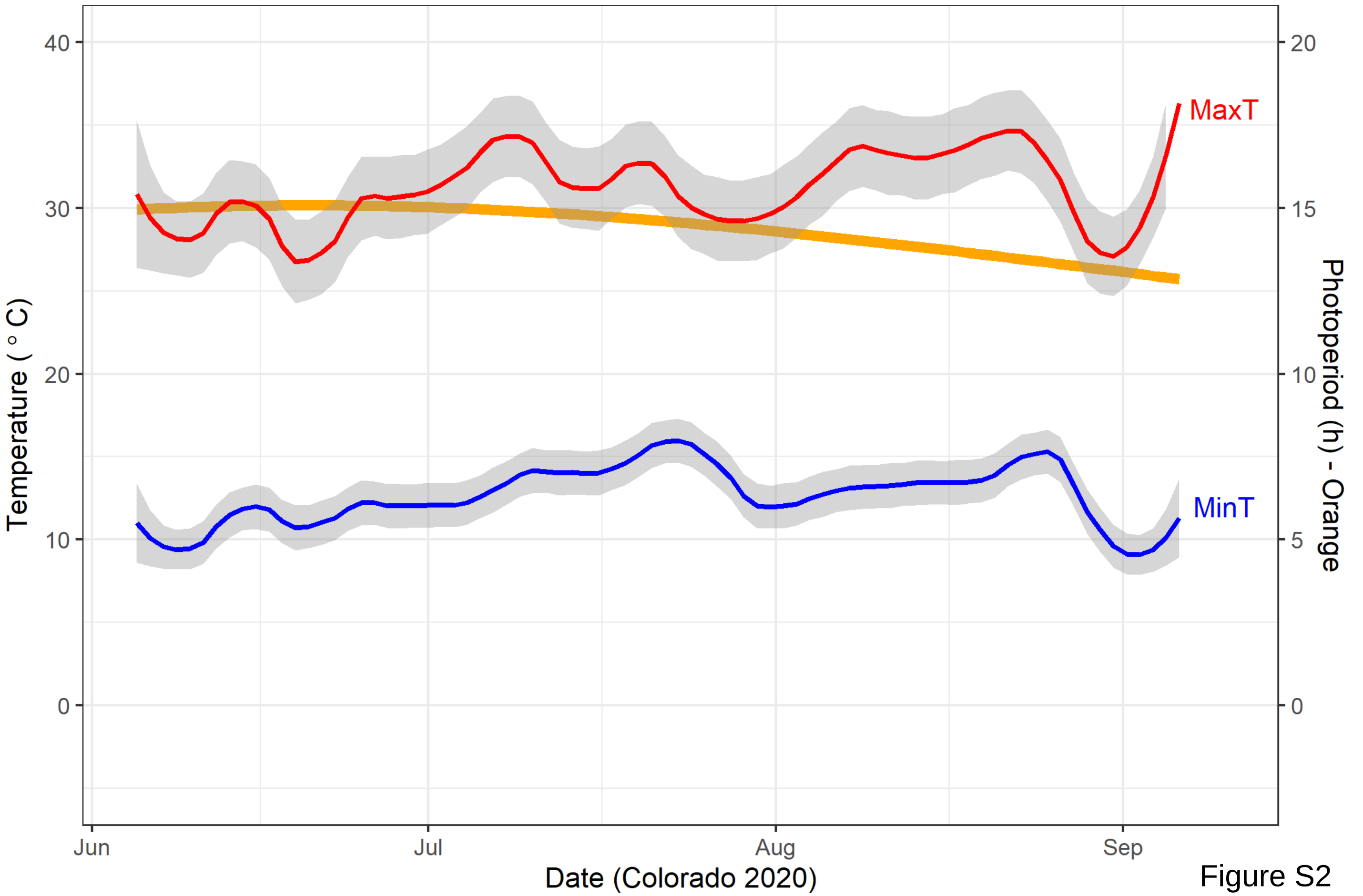

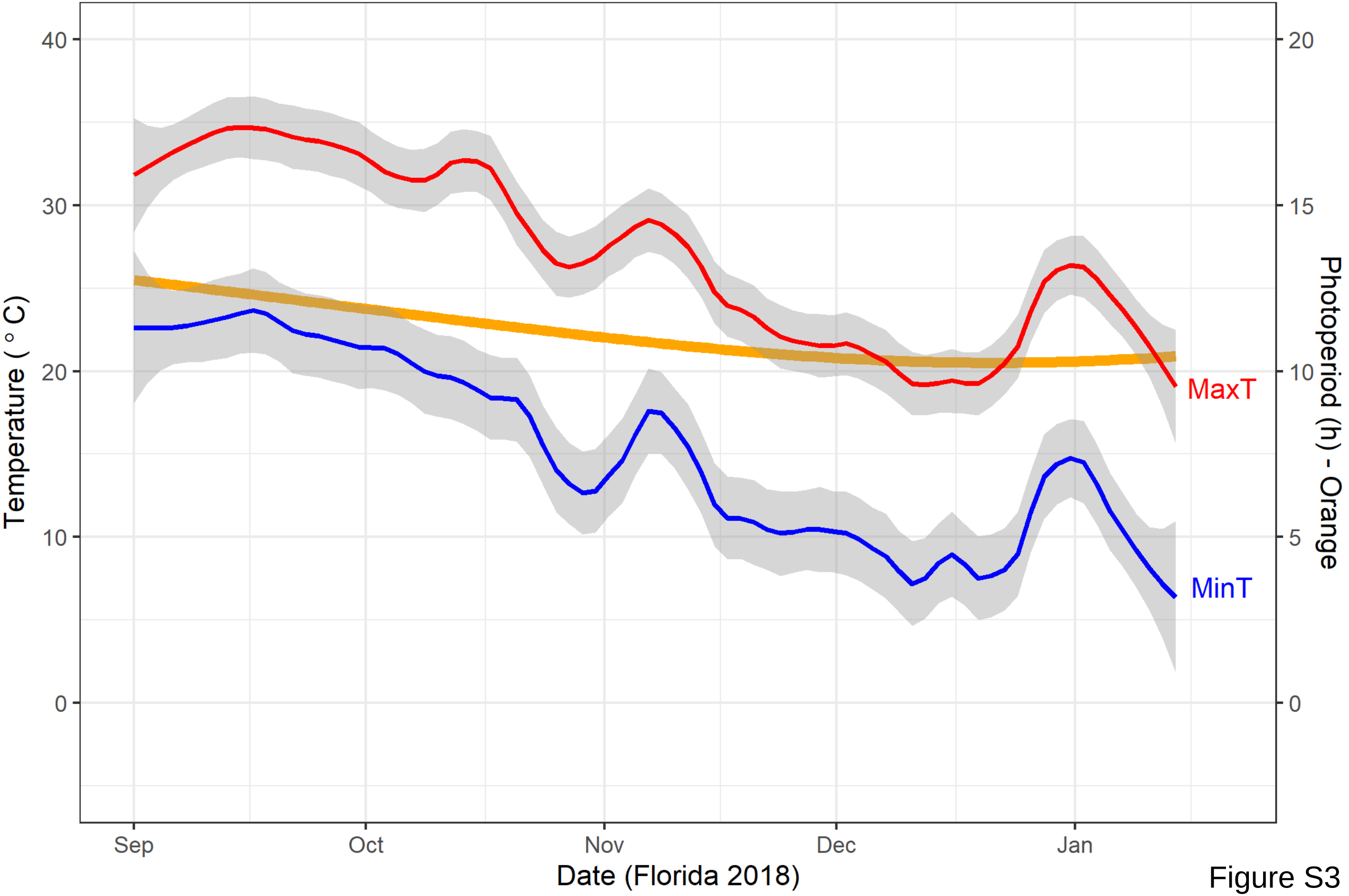

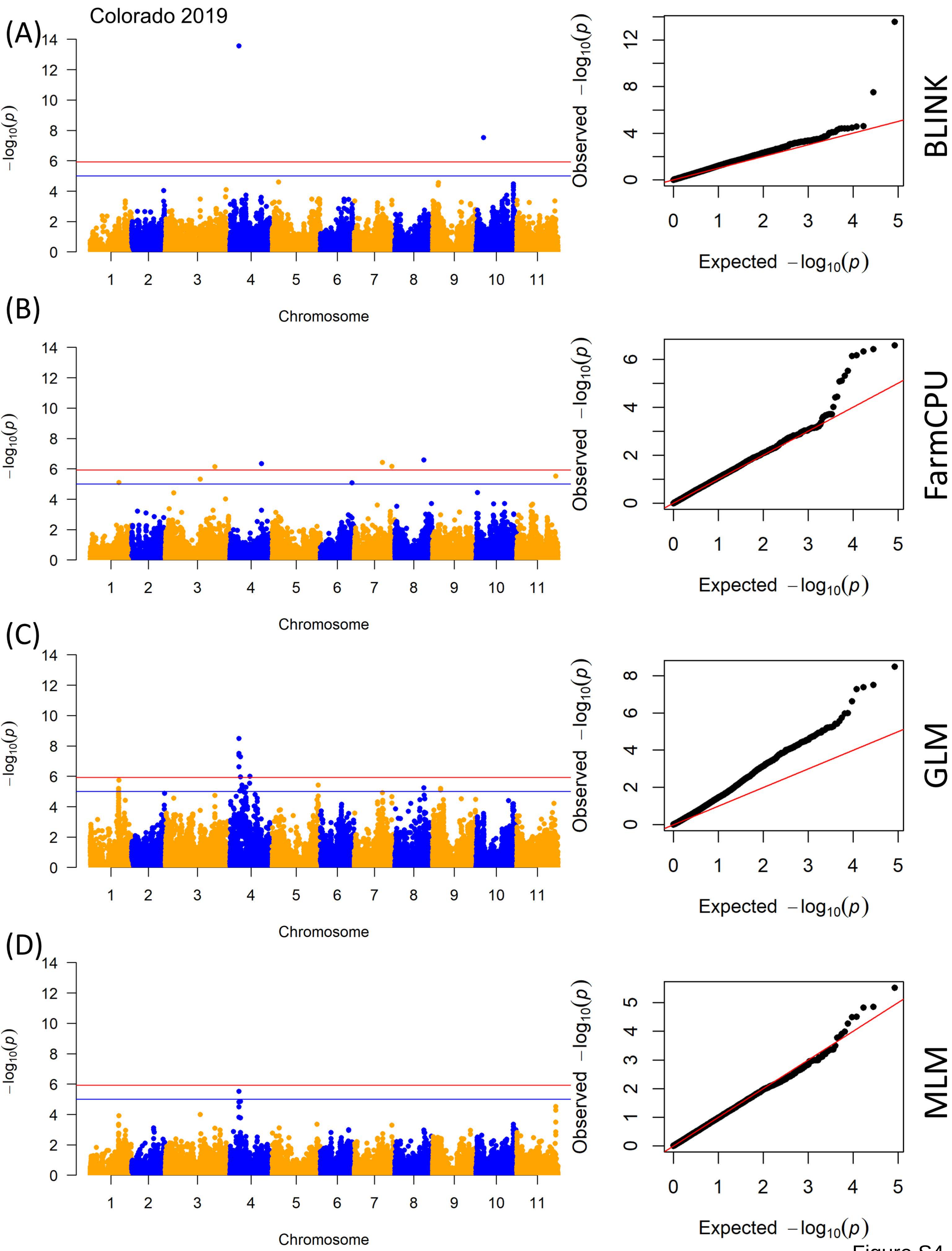

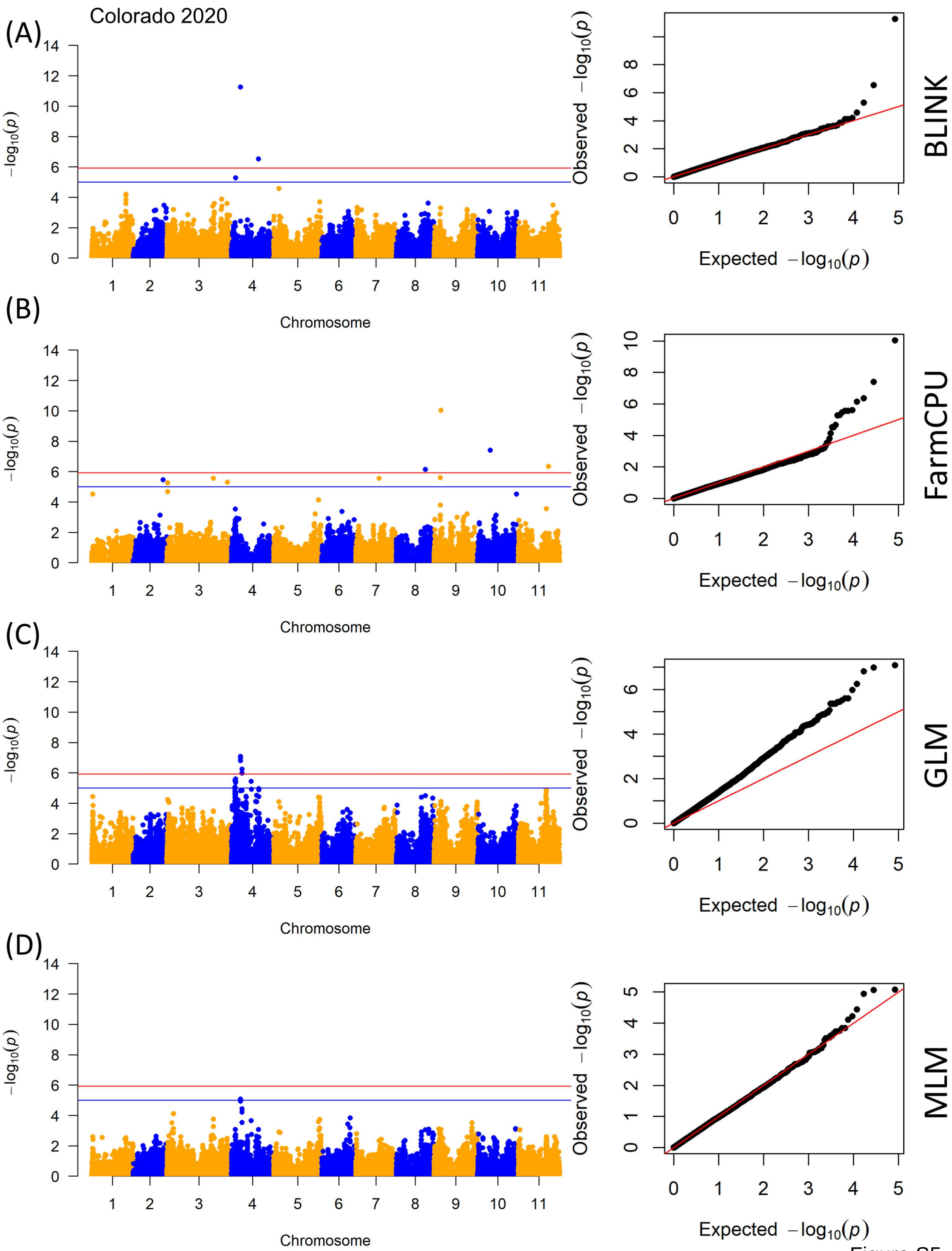

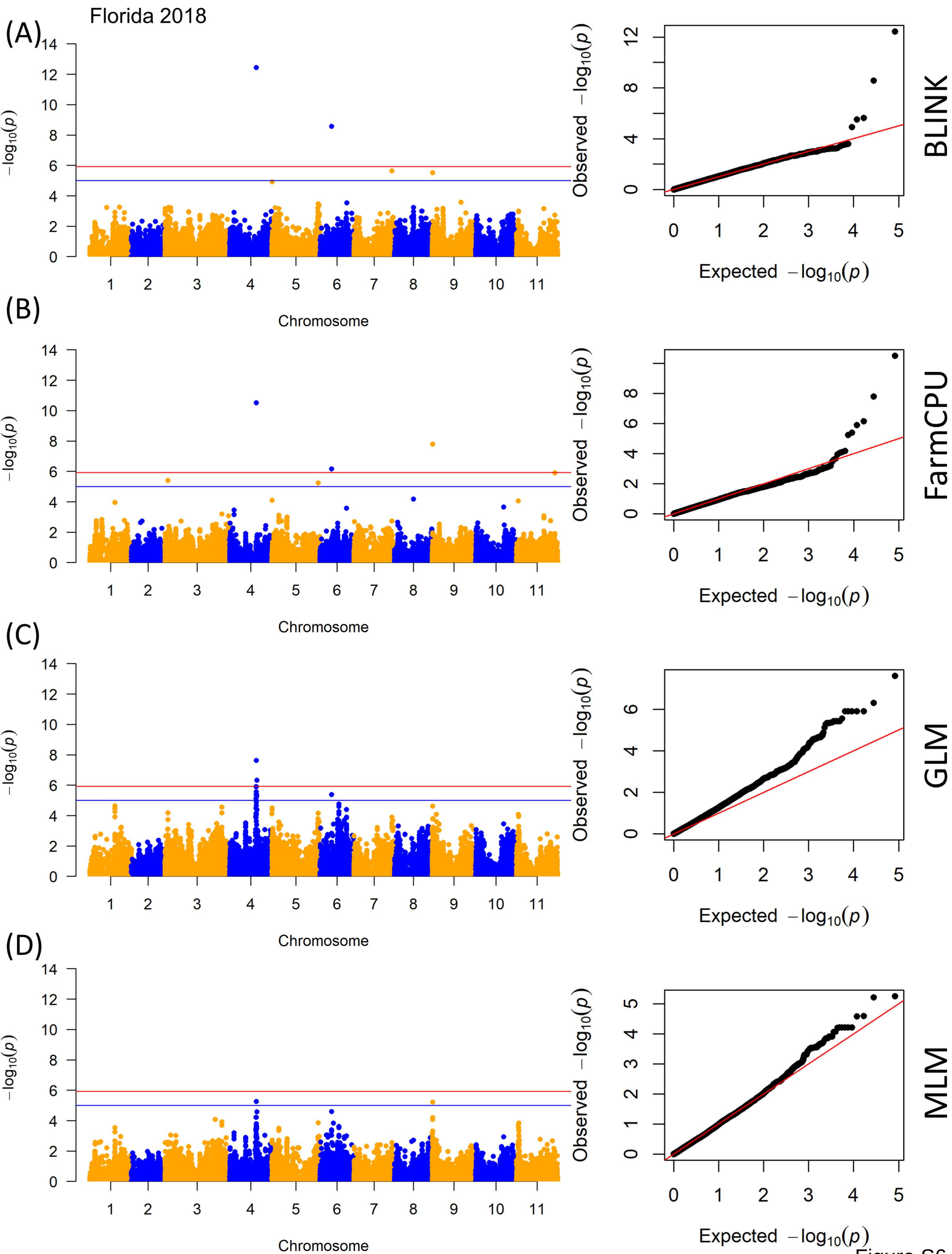
